# mRNA vaccine induces protective immunity against the type III secretory virulence of *Pseudomonas aeruginosa*

**DOI:** 10.1101/2023.06.09.544431

**Authors:** Ken Kawaguchi, Mao Kinoshita, Kazuki Sudo, Keita Inoue, Yoshifumi Naito, Makoto Oba, Satoshi Uchida, Teiji Sawa

## Abstract

Few studies have addressed mRNA vaccination against bacteria despite the desperate need to control antimicrobial resistance. Herein, we developed mRNA vaccines targeting the type III secretion system of *Pseudomonas aeruginosa*, which plays a critical role in the pathogenesis. Vaccination with a nucleoside-modified mRNA encapsulated in lipid nanoparticles improved mouse survival, reduced the bacterial load in the lung, and alleviated pathological changes in the lungs after bacterial challenge.

## Main text

The antimicrobial resistance (AMR) of pathogenic bacteria poses a significant global public health problem.^1^ AMR is expected to become more prevalent and cause an increasing number of deaths in the near future. This emphasises the urgent need for alternative approaches to antibiotics, including preventive vaccines.^2^ *P. aeruginosa* is an important AMR pathogen that causes life-threatening infections, particularly in elderly and immunocompromised patients.^3^ Despite decades of vigorous vaccine development, no licenced vaccines against *P. aeruginosa* are available.^4^ However, recent advances in mRNA vaccines have changed the landscape.^5,6^ Indeed, mRNA-based COVID-19 vaccines have shown robust vaccination efficiency and high scalability to billions of doses without serious safety concerns. The billions of doses administered worldwide provide a wealth of clinical data. In addition, mRNA vaccines can reduce developmental costs due to their versatility in targeting multiple pathogens with a single platform. However, the primary targets of mRNA vaccines are viruses, and only a few reports have addressed their use against bacterial infections.^7,8^ To the best of our knowledge, no previous studies have used mRNA vaccines against *P. aeruginosa*.

The selection of appropriate antigens is critical for vaccines against bacteria, as they possess a much larger number of genes than viruses. Our previous studies identified the PcrV protein at the tip of a type III secretion system (TTSS) as an effective target for active and passive immunisation against *P. aeruginosa*.^*9,10*^ *P. aeruginosa* exerts its pathogenicity by injecting protein toxins into target eukaryotic cells through the TTSS. The phospholipase toxin ExoU, a type III secretory toxin of *P. aeruginosa*, induces acute lung epithelial injury and alveolar macrophage death, and plays a pivotal role in lung injury.^11-13^ A specific antibody against PcrV, which is located as a cap-like structure at the tip of the secretory needle of the TTSS apparatus, can counteract this virulence by inhibiting the translocation of the toxin into eukaryotic cells.^14^ Thus, blocking PcrV alleviates lung damage and improves survival in mice after bacterial challenge. Ultimately, this strategy has demonstrated feasibility in a clinical trial.^15^ These findings led us to design *pcrV* mRNA for mouse vaccination experiments in this study. Using chemically modified mRNA and a lipid nanoparticle (LNP), we provide the proof of concept for using mRNA vaccines to prevent *P. aeruginosa* infection.

First, we evaluated the effect of PcrV antigen secretion from mRNA-introduced cells on antigen production (**Figure 1A**). Mice were vaccinated twice every 3 weeks for serum collection 2 weeks after the second dose. The addition of the human tissue plasminogen activator secretion signal (tPAss) to the *pcrV* mRNA (ss*-pcrV*) markedly increased anti-PcrV IgG production (**Figure 1B**). However, vaccination efficiency was still low in some mice.

**Figure 1.**
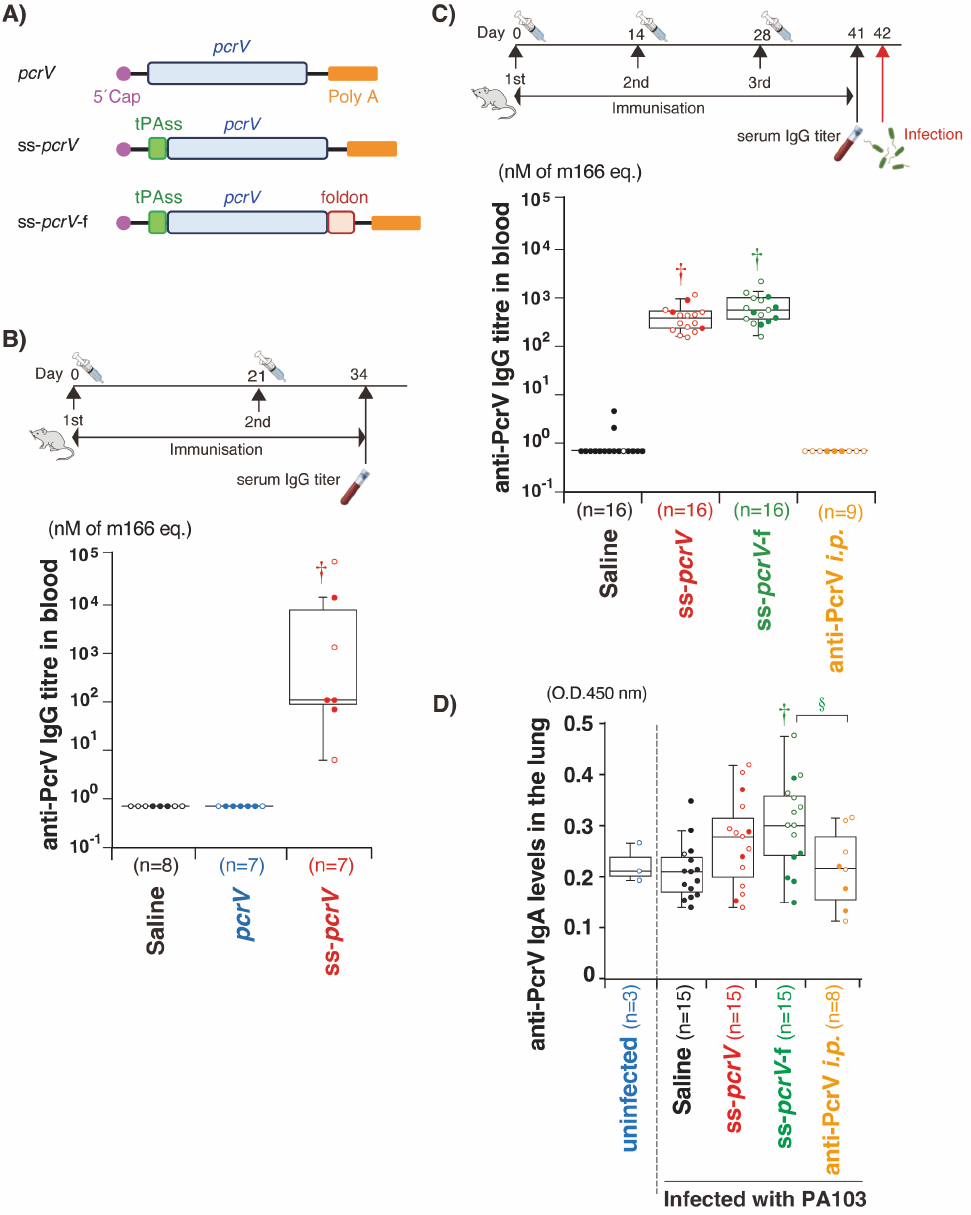
Vaccine designs and antibody titres. (A) Designs of the mRNAs. (B, C) Anti-pcrV IgG titres in mouse serum. (D) Anti-pcrV IgA levels in the lung. †*p* < 0.05 compared with the saline group, §*p* < 0.05 compared with the anti-PcrV i.p. group.

This issue led us to change the vaccination schedule to 3 times every 2 weeks and to add a foldon, a trimerisation domain, at the C-terminus of PcrV in a bacteria challenge experiment (see the upper panel in **Figure 1C** for the experimental schedule). Intraperitoneal injection of the anti-PcrV IgG antibody 1 h before the challenge was performed as a positive control of immunisation. Note that the IgG injection was performed after blood collection for IgG titre measurement in **Figure 1C**, providing no increase in IgG titre in the measurement. The 3-times vaccination protocol resulted in high anti-PcrV IgG titres in all 16 mice injected with ss-*pcrV* mRNA (**Figure 1C**). Adding a foldon to ss-*pcrV* did not further increase the titre. We also quantified IgA levels from homogenates of the lungs obtained one day after the viral challenges, as IgA plays a critical role in mucosal immunity. Intriguingly, vaccinations using ss-*pcrV* and ss-*pcrV*-f mRNA tend to increase IgA levels with the ss-*pcrV*-f mRNA group showing statistical significance compared to saline-treated and IgG-injected groups (**Figure 1D**). From a safety viewpoint, LNP injection induced only minimal changes in blood tests (**Supplementary Table S1**).

On day 42 after the first immunisation, the mice were challenged intratracheally with *P. aeruginosa* PA103 (1.0 × 10^6^ colony-forming units [CFU]). All infected mice showed a drastic drop in body temperature at 4 h, indicating successful infection with PA103 (**Figure 2A**). More precisely, the body temperatures of the ss-*pcrV* and ss-*pcrV*-f groups were significantly higher than those of the saline-treated group 4 and 8 h after the challenge. Remarkably, the survival rate was significantly higher for mice treated with ss-*pcrV-* and ss-*pcrV*-f mRNA (75.0% and 62.5%, respectively) than those in the saline-injected infected group (6.25%) 24 h after the challenge (**Figure 2B**). Vaccination with ss-*pcrV* and ss-*pcrV*-f mRNA also reduced the total lung bacterial counts by 10–30-fold compared with the counts in the saline-treated group in surviving and dead mice 24 h after the challenge (**Figure 2C**). Furthermore, the inflammatory response in the lungs, another critical inducer of lung damage, became milder after vaccination with ss-*pcrV* and ss-*pcrV*-f mRNA, with decreased myeloperoxidase activity and interleukin 6 (IL-6) levels in the lungs compared to the saline-treated group (**Figure 2D**,**E**). Finally, histological analyses at 24 h post-challenge supported these findings (**Figure 2F-J**). Saline-treated control mice showed enhanced neutrophil infiltration, alveolar haemorrhage, and destruction of the alveolar structure (**Figure 2G**). In contrast, these pathological changes were minimal in mice vaccinated with ss-*pcrV* or ss-*pcrV*-f mRNA (**Figure 2H, I**). Meanwhile, adding a foldon domain to the pcrV antigen did not provide an additional vaccination effect compared to ss-*pcrV* mRNA, as shown by the IgG titre measurement (**Figure 1C**).

**Figure 2.**
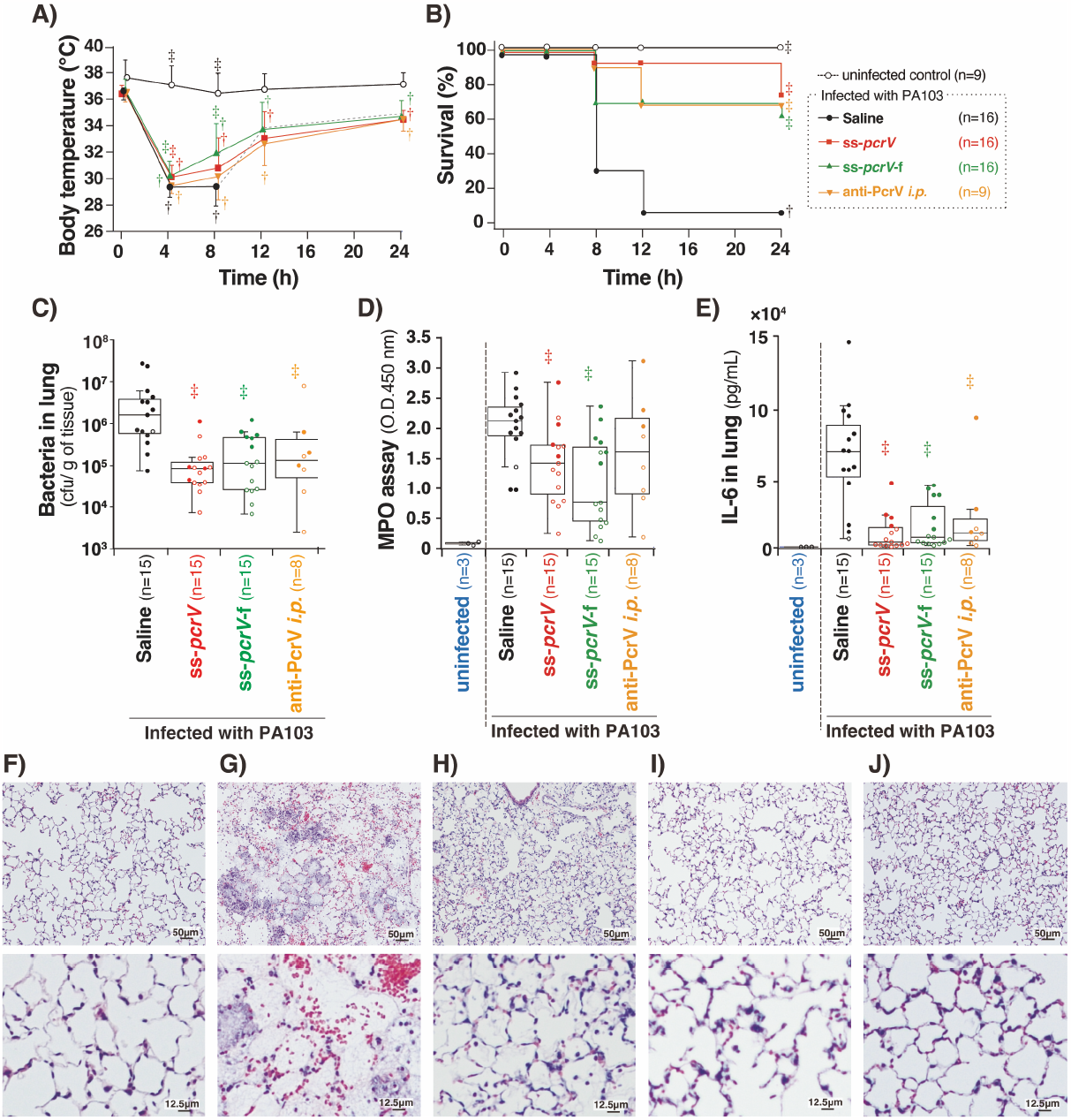
Vaccination effects against bacterial challenge. (A) Body temperature. (B) Survival. (C) Bacterial load, (D) myeloperoxidase activity, (E) interleukin 6 levels in the lungs, and (F-J) histological changes 24 h after the challenge. (F) An uninfected control. (G) Infected mice treated with saline. (H) ss-*pcrV* mRNA. (I) ss-*pcrV*-f mRNA. (I) nti-PcrV antibody (J). See **Figure 1C** for the experimental schedule. †*p* < 0.05 compared with the uninfected control group. ‡*p* < 0.05 compared with the saline-injected PA103-infected group.

The present study is the first to demonstrate the utility of mRNA vaccines against *P. aeruginosa* infections. Vaccinations alleviated lung injury presumably by inhibiting the injection of the toxin into lung cells by the TTSS (**Figure 2F-J**). However, this mechanism apparently does not explain the effect of vaccination on bacterial growth in the lungs (**Figure 2C**). According to our previous study, anti-PcrV antibodies improve the survival of alveolar macrophages by preventing TTSS-mediated toxic effects on macrophages.^9^ Therefore, the mRNA-based immunisation used in the present study may have reduced the bacterial burden by preserving macrophage function, thus enabling them to kill the bacteria in the lungs. Furthermore, the mRNA vaccines alleviated inflammation, which is another inducer of lung damage (**Figure 2D**,**E**). These mechanisms may contribute to the improved mouse survival rates. These results are remarkable as post-translational modifications of PcrV expressed from mRNA in mammalian cells may differ from those of PcrV expressed in bacteria, potentially reducing the antigenicity of PcrV.^16^ In contrast, when targeting viruses, major targets of mRNA vaccines, this issue is not relevant, as viruses produce antigens via the host mammalian machinery. Robust immunisation with mRNA vaccines may override species differences in post-translational modifications during vaccination against bacteria.

*P. aeruginosa* is not a supervirulent pathogen, but rather, an attenuated pathogen that causes diseases in vulnerable patients with special conditions, such as immunodeficiency and pulmonary cystic fibrosis.^3^ This complicates the acceptable development costs, the selection of target patients, and the implementation of clinical trials. The scalability and cost-effectiveness of mRNA vaccines offer a promising solution to this problem. Another advantage of mRNA vaccines is the flexibility to change antigens simply by redesigning the mRNA sequences. In this regard, many pathogenic gram-negative bacteria share the TTSS as a highly homologous virulence factor, providing a potential target for mRNA vaccines by changing RNA sequences.^17^ Therefore, the potential applications of this mRNA-based approach are broad.

## Materials and Methods

### Preparation of mRNA

Plasmid DNA (pDNA) possessing the pcrV coding sequence under the control of the T7 promoter, with or without tPAss sequences (MDAMKRGLCCVLLLCGAVFVSAR) and foldon sequences (GYIPEAPRDGQAYVRKDGEWVLLSTFL), was constructed by GenScript Japan (Tokyo, Japan). Template DNA was prepared by polymerase chain reaction (PCR) amplification of pDNA, using a reverse primer containing an 80-nt poly T sequence to add a poly A tail to the mRNA. For *in vitro* transcription, an anti-reverse capping analogue (Trilink Biotechnologies, San Diego, CA, USA), N1-methylpseudouridine (Sapala Organics Private Limited, Telangana, India), and a MegaScript T7 transcription kit (Thermo Fisher Scientific, Waltham, MA, USA) were used. LNPs from ALC-0315 (MedChemExpress, Monmouth Junction, NJ, USA), ALC-0159 (MedChemExpress), 1,2-distearoyl-sn-glycero-3-phosphocholine (Fujifilm Wako, Osaka, Japan), and cholesterol (Sigma-Aldrich, St. Louis, MO, USA) were prepared with a nitrogen/phosphate ratio of 6, as previously reported.^18^ The prepared LNPs had average sizes of less than 100 nm and narrow polydispersity indices of approximately 0.15 **(Supplementary Figure S2**).

### Recombinant PcrV and anti-PcrV antibodies

Endotoxin-free recombinant PcrV (rePcrV) was prepared as reported previously^19^. Briefly, the coding sequence for PcrV was amplified from the chromosome of *P. aeruginosa* PAO1 by PCR. The PCR fragments were ligated into an *Escherichia coli* expression vector (pQE30; Qiagen, Hilden, Germany) to create a protein construct with a six-tandem histidine residue (His-tag) at the amino-terminus. The expression vector was introduced into *E. coli* strain M15 (Qiagen), and recombinant proteins were produced from the *E. coli* culture and induced by isopropylthio-β-galactoside. Crude proteins were purified using nickel-nitrilotriacetic acid agarose columns (Novex R901-15, Thermo Fisher Scientific), dialysed against phosphate-buffered saline (P3813, Sigma-Aldrich) for 72 h, and applied to endotoxin removal resin columns (Pierce, Thermo Fisher Scientific). The purity of rePcrV was evaluated by sodium dodecyl sulphate-polyacrylamide gel electrophoresis, after which an intense single band appeared on the stained gel.

The anti-PcrV polyclonal IgG fraction was prepared from a re-PcrV-vaccinated rabbit (Kitayama Labs, Ina, Japan), as previously reported.^19^ Murine monoclonal anti-PcrV IgG^2b^ m166^20,21^ was purchased from Creative Biolabs (New York, NY, USA).

### Animals

Certified pathogen-free male ICR mice (4 weeks old; body weight, 20–25 g) were obtained from Shimizu Laboratory Supplies Co. Ltd. (Kyoto, Japan). The mice were housed in cages (with filter tops) under pathogen-free conditions. The protocols for all animal experiments were approved by the Animal Research Committee of the Kyoto Prefectural University of Medicine (approval number: M2022-509) prior to starting the experiments.

### Vaccination

mRNA-LNPs were dissolved in a 70 μL solution and intramuscularly injected into mice at a dose of 10 μg/mouse. The vaccination schedules are shown in **Figure 1B** and **C**. For antibody titre measurements, approximately 50 μL of peripheral blood was collected from the tail vein using a haematocrit capillary tube. One group of mice (anti-PcrV group) intraperitoneally received 100 μL of ovine anti-PcrV polyclonal IgG 1 h before the infection challenge. Note that this passive immunisation has been shown to exert a protective effect against lethal *P. aeruginosa* infection. ^*9,22*^

### Infectious challenge by *P. aeruginosa*

The *P. aeruginosa* PA103 strain, originally isolated from a patient in Australia in the late 1960s, was used in the challenge assay.^23^ This strain has a cytotoxic phenotype, with positive type III secretion of ExoT and ExoU, and it carries the type III secretion system genotype *exoS*^−^, *exoT*^+^, *exoU*^+^, and *exoY*^+^. However, it exhibits an ExoY secretion-negative phenotype caused by a codon mutation. Bacterial suspensions were prepared as described previously.^19^ The cultures were grown at 32°C for 12 h in a shaking incubator and centrifuged at 3,000 rpm for 10 min. The bacterial pellet was washed thrice in saline and diluted to the appropriate number of CFU per mL, as determined by spectrophotometry. The number of bacteria was determined by assessing the CFU of the diluted aliquot on a sheep blood agar plate. The solution containing PA103 (1.0 × 10^6^ CFU in 60 μL of saline) was instilled into the lungs of each vaccinated mouse through an endotracheal needle, as described previously, under light inhalational anaesthesia using sevoflurane.^24^ The mice in the saline (no infection) group were intratracheally administered saline. The survival and body temperature of all mice were monitored for 24 h, after which the survivors were euthanised, and the lungs of each mouse were collected, weighed, and homogenised for further evaluation.

### IgA quantification, myeloperoxidase (MPO) activity, cytokine levels, and bacteriological assay

Wet lung weights were measured as an index of lung oedema at the 24-h time point, as described previously.^25^ IgA levels in lung homogenates were measured by ELISA with goat anti-mouse IgA a chain antibody conjugated to horseradish peroxidase (ab97235; Abcam, Cambridge, UK), as previously reported.^*10*^ MPO activity was measured in lung homogenates as reported previously^25^. Interleukin-1β, IL-6, and tumour necrosis factor α concentrations were quantified in lung homogenates using an ELISA kit (BD Opt EIA Mouse IL-6 ELISA Sets, #555240; BD Biosciences Franklin Lakes, NJ, USA) according to the manufacturer’s instructions. The sequentially diluted lung homogenate was inoculated onto a sheep blood agar plate and incubated at 37°C overnight to calculate the number of remaining bacteria per gram of lung tissue.

### Histopathological assay

A designated mouse from each group was euthanised for histological analysis 24 h after infection. The lungs were perfused with a 10% formalin neutral buffer solution for fixation and embedded in paraffin. Sections were stained with haematoxylin and eosin.

### Statistical analysis

For multiple comparisons of survival curves, post-hoc analysis for the log-rank test was used with *p*-values adjusted by the Benjamini and Hochberg method using RStudio (2022.7.1. Building 554, R version 4.2.1, R Studio PBC). One-way analysis of variance and unpaired Student’s t-tests were used to analyse body temperature. The antibody titre, lung weight, distance travelled, and number of bacteria in the lungs were evaluated using the Kruskal-Wallis test. A *p*-value less than 0.05 was considered statistically significant. KaleidaGraph (Ver.5.0.2; Synergy Software, Reading, PA, USA) was used to visualise the data as graphs. Body temperature data are expressed as the mean ± standard deviation, and other data are expressed as the median (interquartile range).

## Author Contributions

Conceptualization, S.U. and T.S.; methodology, S.U. and T.S.; investigation, K.K., M.K., K.S., and Y.N.; writing—original draft preparation, S.U. and T.S., K.I. and K.Mo.; writing—review and editing, S.U. and T.S.; supervision, S.U. and T.S.; project administration, S.U. and T.S.; funding acquisition, S.U. and T.S. All authors have read and agreed to the published version of the manuscript.

## Funding

This work was supported by Leading Advanced Projects for medical innovation: LEAP (No. 21gm0010008h0001, P.I.: Hiroshi Abe, Graduate School of Science, Nagoya University) to U.S. and T.S. and Research Program on Emerging and Re-emerging Infectious Diseases (No. 21fk0108620h0001) to S.U. from Japan Agency for Medical Research and Development and the Japan Society for the Promotion of Science, Grant-in-Aid for Scientific Research KAKENHI from the Ministry of Education, Culture, Sports, Science and Technology, Japan (Nos. 24390403, 26670791, 15H05008, 18H02905, and 22H03176)to T.S. and (No. 21H04962) to U.S.

## Data Availability Statement

The datasets generated and/or analyzed in this study are available from the corresponding author upon reasonable request.

## Acknowledgments

We thank Sachi Ibuki (Kyoto Prefectural University of Medicine), Erika Mochizuki, Nao Horii (Tokyo Medical and Dental University) and Miki Masai (NanoCarrier Co.) for technical assistance.

## Conflicts of Interest

S.U is a founder of Crafton Biotechnology.

