## Supplementary information for "mRNA vaccine induces protective immunity against the type III secretory virulence of *Pseudomonas aeruginosa*"

**Supplementary Table S1** Blood chemistry 24 h after the vaccination.

| | AST (IU/L) | ALT (IU/L) | LDH (IU/L) | BUN (mg/dL) | Cre (mg/dL) |
|---|---|---|---|---|---|
| Untreated | 51.8 ± 4.4 | 44.4 ± 5.7 | 139 ± 26 | 23.2 ± 1.4 | 0.1 ± 0.0 |
| LNP | 83.4 ± 20.0 | 53.0 ± 9.4 | 275 ± 92 | 26.9 ± 1.3 | 0.2 ± 0.0 |

Measured by Dri-Chem 7000IZ system (FujiFilm, Tokyo, Japan). *n* = 5. Data are shown as the mean ± standard deviation.
Abr.: AST, aspartate aminotransferase; ALT, alanine aminotransferase; LDH, lactate dehydrogenase; BUN, blood urea nitrogen; Cre, Creatinine

**Supplementary Table S2** Dynamic light scattering (DLS) measurement of LNPs.

| | Size (d, nm) | PDI |
|---|---|---|
| LNP (ss-pcrV) | 83.4 ± 1.0 | 0.150 ± 0.009 |
| LNP (ss-pcrV-f) | 85.5 ± 1.2 | 0.141 ± 0.008 |

Measured using Zetasizer Nano ZS (Malvern, Herrenberg, Germany). Three measurements were performed using LNPs prepared separately for each time. Data are shown as the mean ± standard deviation.
